# Multiparametric Signature of Glioblastoma Differentiation Revealed by Imaging of Cellular Epigenetic Landscapes

**DOI:** 10.1101/348888

**Authors:** Chen Farhy, Santosh Hariharan, Jarkko Ylanko, Lotte van Woudenberg, Flavio Cimadamore, Fernando Ugarte, Camilla Forsberg, Chun-Teng Huang, David W. Andrews, Alexey V. Terskikh

**Affiliations:** Sanford Burnham Prebys Medical Discovery Institute, La Jolla, California, USA; Biological Sciences Platform, Sunnybrook Research Institute and Department of Medical Biophysics, University of Toronto, Ontario, Canada; Department of Biomolecular Engineering, Institute for the Biology of Stem Cells, University of California, Santa Cruz, California, USA; Department of Biochemistry, University of Toronto, Ontario, Canada

## Abstract

The resistance of Glioblastoma (GBM) to conventional cytotoxic drugs has prompted novel therapeutic strategies, including differentiating tumor propagating cells (TPCs) into less tumorigenic cells using small molecule inducers of TPC differentiation. However, high-throughput screening for such molecules is hampered by the lack of robust markers of GBM differentiation. To obtain a signature of differentiated TPCs, we developed “Microscopic Imaging of Epigenetic Landscapes” (MIEL), which captures patterns of nuclear staining for epigenetic marks to derive feature-fingerprints of individual cells. We confirmed MIEL’s ability to accurately distinguish multiple cell fates and identified a multiparametric epigenetic signature of differentiated TPCs. Critically, we validated epigenetic imaging-based signature using global gene expression thus providing the proof of principle for the MIEL’s ability to select and prioritize small molecules, which induce TPC differentiation.

## Introduction

Malignant GBM is the most common and lethal brain tumor, however, current therapeutic options offer little prognostic improvement, and the median survival time has remained virtually unchanged for several decades (1–3). GBM tumor mass is a heterogeneous mix of cells expressing lineage markers found in neural stem/precursor cells, neurons, and glia, and aberrantly expressed proliferation markers (4,5). TPCs are a subpopulation of GBM cells with increased tumorigenic capability (6) operationally defined as early passaged (<15) GBM cells propagated in serum-free medium (7).

Compared to the bulk of the tumor, TPCs are more resistant to drugs, such as temozolomide (TMZ), and radiation therapy (8,9). This resistance may explain the failure of traditional therapeutic strategies based on cytotoxic drugs targeting GBM. A promising alternative approach aims to drive differentiation of tumor cells, particularly TPCs, thereby reducing tumor expansion through decreased cell proliferation and increasing sensitivity to cytotoxic treatments (10–15).

Culturing primary GBM cells in serum-containing medium induces their differentiation into cells with drastically reduced tumorigenic potential (16). In addition, Bone Morphogenetic Protein 4 (BMP4) treatment was reported to induce GBM differentiation (17,18), which might be reversible (19) and is contingent on the presence of functional BMP receptors (20). These observations support the potential therapeutic value of small molecules that mimic the differentiation effect of serum and BMPs on TPCs. Several studies successfully performed high content screenings using normal neural progenitor and monolayer cultures of GBM to identify cytotoxic molecules (21,22). However, attempts to design screening strategies identifying inducers of GBM differentiation have been met with multiple difficulties. One critical problem is the lack of informative markers for GBM differentiation. The most commonly used markers (e.g. sex-determining region Y-box 2 (SOX2) and glial fibrillary acidic protein (GFAP)), exhibit highly variable expression in TPCs (4) and in our hands these markers failed to prioritize molecules mimicking serum/BMP ability to differentiated GBM. In addition, recent single-cell expression analysis of primary GBM was unable to identify a limited set of GBM differentiation markers that could be used for high-throughput screening (23).

Here, we introduce MIEL, a novel phenotypic screening platform that takes advantage of epigenetic modifications and multiparametric image analysis to reveal a signature of TPC differentiation amenable to high-content screening. We have validated MIEL’s ability to select and prioritize small molecules mimicking serum/BMP4 effect on GBM using global gene expression profiling. This new approach opens the door for discovering small molecule drugs that can phenocopy the effect of biologicals such as serum or BMP known to induce GBM differentiation.

## Results

### Brief treatment with serum or Bmp4 initiates TPC differentiation

A comparative analysis of gene expression changes in TPCs following short serum or Bmp4 treatment, which is relevant to our high-throughput screening objective, has not been conducted. We therefore treated several GBM cell lines for 3 days with serum or Bmp4 and then quantified expression of core transcription factors previously shown to determine the transcriptomic program of TPCs (6). Immunostaining revealed that the 4 transcription factors Sox2, Sall2, Brn2 and Olig2 were down regulated by both serum and Bmp4 in a cell line dependent manner (Supplementary Fig. 1a). After 3 days of treatment, the growth rate of TPCs was reduced by both serum and Bmp4 (Supplementary Fig. 1b). RNAseq analysis of serum and Bmp4 treated GBM2 cells revealed that 3 days treatment reduced (vs untreated cells) the expression of most genes previously found to constitute the transcriptomic stemness signature (23) (Supplementary Fig. 1c). To identify the cellular processes altered by these treatments, we conducted differential expression analysis. We found that expression of 4852 genes was significantly altered (p<0.01 and -1.5<Fold Change >1.5) by either serum or Bmp4 treatment. Gene Ontology (GO) analysis of these altered genes indicated enrichment in multiple GO categories consistent with initiation of TPC differentiation – including cell cycle, cellular morphogenesis associated with differentiation, differentiation in neuronal lineages, histone modification, and chromatin organization (Supplementary Fig. 2). Taken together, these results demonstrate that a 3 days treatment with serum or Bmp4 is sufficient to result in transcriptomic changes characteristic of TPC differentiation.

### Sox2- and GFAP-based screening doesn’t prioritize inducers of TPC differentiation

Several studies suggest that Sox2 function is required for maintenance of TPCs and that its knockdown induces TPC differentiation (6,15,24,25). We therefore selected Sox2 as a marker of the TPC state. For the differentiated state, we selected GFAP, an astrocytic marker previously shown to be upregulated following differentiation of TPCs (6,24). We confirmed that Bmp4, but not serum, treatment also increased GFAP expression (Supplementary Fig. 1a).

Among several GBMs tested, the GBM2 line exhibited the largest reduction in Sox2 and increase in GFAP and was selected for screening. GBM2 TPCs were plated in 384-well plates, treated with the Prestwick library compounds (10 μM, 1200 molecules) for 3 days, fixed, and then immunostained for Sox2 and GFAP. Hits were defined as compounds that increased GFAP and decrease Sox2 by more than 40% or any compounds that decrease Sox2 alone by more than 100%; Bmp4 was used as a positive control (Fig. 1a). We detected 19 hits (Supplementary Fig. 3), however, reduced cell viability and the presence of pyknotic nuclei indicated apparent cytotoxicity of the most hits (z-score for viable cell count less than -4 compared with -2.33 for Bmp4; Supplementary Fig. 3). We therefore retested the hit compounds at lower concentrations (3, 1 and 0.3μM) and observed that a 3 days treatment with 0.3μM Digitoxigenin, a Na^+^/K^+^ ATPase inhibitor, was able reduce Sox2 expression while maintaining growth rate and Ki67 expression levels similar to Bmp4 (Fig. 1b, c and supplementary Fig. 10a). A related compound Digoxin also reduced Sox2 expression but induced a stronger reduction in growth rate (Fig. 1b, c and supplementary Fig. 10a); both Digitoxigenin and Digoxin were able to downregulate expression of the core transcription factors similar to serum and Bmp4 (Fig. 1c). To validate these results using a “gold standard” of cell fate analysis, we conducted a whole genome expression analysis of treated GBM2 TPCs. To test whether Digitoxigenin and Digoxin increased transcriptomic similarity of treated cells to serum or Bmp4 treated cells we used FPKM values of all expressed genes (FPKM>1) to calculate the Euclidean distance between drug and serum or Bmp4 treated cells. We discovered that neither Digitoxigenin nor Digoxin reduced the distance of treated cells to the desired state (Fig. 1d). Several representative GO terms illustrate the gene expression changes induced by digoxin and digitoxigenin which were markedly different from those induced by serum or BMP4 (Fig. 1e). We concluded that our 2 top hits didn’t induce desirable TPC fate change, despite downregulation of some core transcription factors essential for TPC propagation. Our results emphasize the need of developing novel approaches to interrogate TPC differentiation, which are compatible with the high-throughput screening and align well with the entire transcriptome analysis.

**Figure 1.**
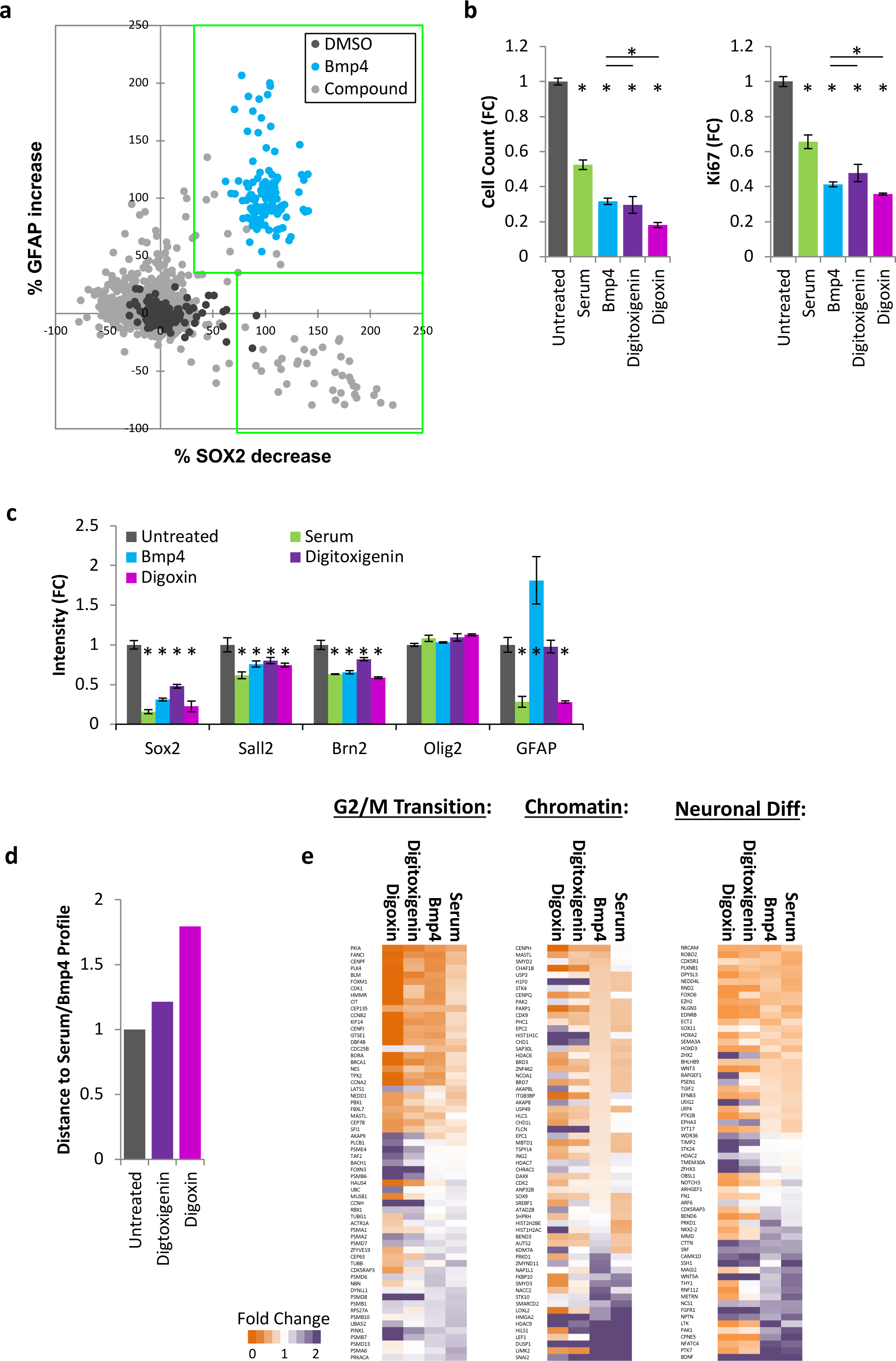
Digoxin and digitoxigenin reduce expression of transcription factors regulating the TPC transcriptomic program but fail to induce GBM differentiation. (a) GBM2 cells were treated with DMSO, Bmp4 or Prestwick compounds (10 μM); after 3 days, cells were immunostained for Sox2 and GFAP. Scatter plot shows % of Sox2 inhibition and % of GFAP activation for individual wells. Compounds showing GFAP>40% and Sox2>40% (large green box) or only Sox2>100% (small green box) were considered hits. (b,c,d) GBM2 cells were treated for 3 days with serum, Bmp4, 0.3 μM digitoxigenin, or 0.3 μM digoxin. Cells were either (b,c) immunolabeled with cell count and intensity normalized to control cells (mean ± S.D, p<0.05, n=3 technical repeats, unpaired two-tailed t-test). (d) Bar graph showing Euclidean distance to serum- or Bmp4-treated cells, calculated using normalized FPKM values of expressed genes for untreated, 0.3μM Digitoxigening or 0.3μM Digoxin treated GBM2 cells. (e) Heat maps showing fold change (RNA sequencing) in expression of select genes taken from the GO list and belonging to 1 of 3 functional classes: cell cycle G2/M phase transition (GO:0044839), chromatin-modification (GO:0006325), and regulation of neuron differentiation (GO:0045664).

### Development of MIEL platform

We developed novel phenotypic screening platform, which interrogates the epigenetic landscape at single cell level using imaged-based machine learning. MIEL takes advantage of epigenetic marks such as histone methylation and acetylation, which are always present in eukaryotic nuclei and can be revealed by immunostaining. MIEL analyzes the immunolabeling patterns of epigenetic marks at the single-cell level – using conventional image analysis methods for segmentation of nuclei, feature extraction and previously described machine learning algorithms (26) (Fig. 2a, Supplementary Fig. 4a, b, and Methods).

**Figure 2.**
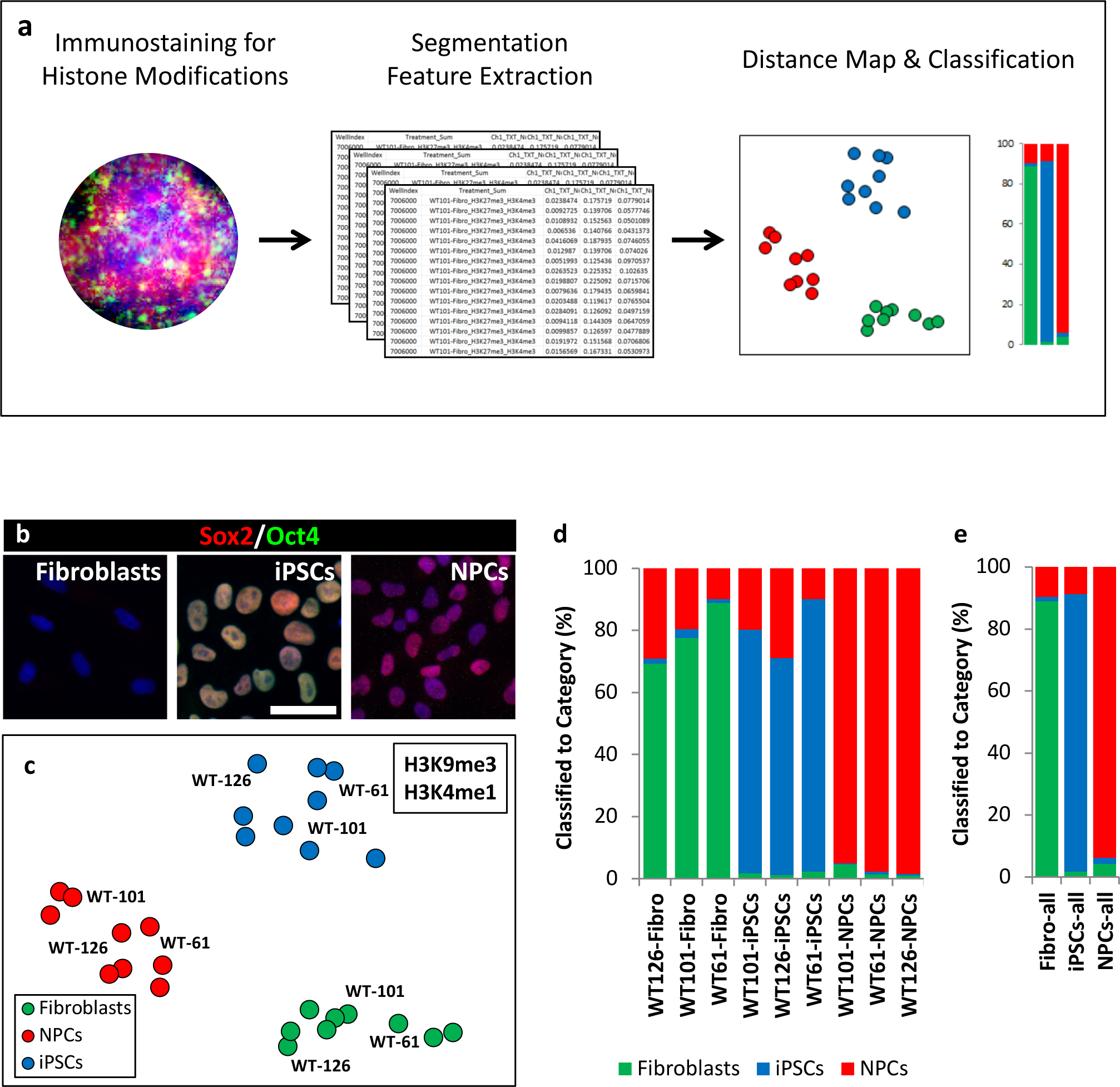
Validation of MIEL platform. (a) Flowchart of MIEL pipeline. Fixed cells were immunostained for the desired epigenetic modifications, stained with Hoechst 33342 to visualize DNA and imaged. Nuclei were segmented based on DNA staining, and texture features calculated from the pattern of immunofluorescence. The relative similarity of multiple cell populations was assessed by calculating the multiparametric Euclidean distance between their pairwise centers and represented in 2D following MDS (distance map). A support vector machine, trained on a random subset of cell data, was used to classify the rest of the individual cells (bar graph). (b) Hoechst 33342 stained (blue), and Sox2 (red) and Oct4 (green) immunofluorescence labeled fibroblasts (Sox2^-^/Oct4^-^), iPSCs (Sox2^+^/Oct4^+^) and NPCs (Sox2^+^/Oct4^-^). Scale bar, 50 μm. (c) Distance map depicting the relative Euclidean distance following MDS between the multiparametric centroids of 9 cell lines: 3 fibroblasts, 3 iPSCs and 3 NPCs, calculated from texture feature values derived from images of H3K9me3 and H3K4me1 marks. Each cell line appears as technical triplicates (ie, dots correspond to images from different wells). (d) Three-way classification of fibroblasts, iPSCs and NPCs from 3 donors. (e) Three-way classification of fibroblasts and NPCs, each pooled from the 3 donors.

Primarily, we utilized 4 histone modifications: H3K27me3 and H3K9me3, which are associated with condensed (closed) facultative and constitutive heterochromatin, respectively; H3K27ac, associated with transcriptionally active (open) areas of chromatin, especially at promoter and enhancer regions; and H3K4me1, associated with enhancers and other chromatin regions (27,28). To focus the learning algorithm on the intrinsic pattern of epigenetic marks, we discarded the intensity and nuclear morphology features and used only texture-associated features (e.g. Haralick’s texture features (29), threshold adjacency statistics, and radial features (30)) for multivariate analysis. Previous studies have successfully employed similar features for cell painting techniques combined with multivariate analyses to accurately classify subcellular localization of proteins (30), cellular subpopulations(31), and drug mechanisms of action (26,32–34). We interpreted the observed patterns as a 2D projection of the 3D topological distribution of a given epigenetic mark in the nucleus. Although this representation degrades the spatial information, the resulting 2D textures, such as foci of high and low intensity, are visually apparent in the computer-enhanced images (Supplementary Fig. 4a).

### MIEL analysis provides signatures of cell fates

We developed MIEL to distinguish between differentiated and undifferentiated TPCs, and to obtain a multiparametric signature of differentiated TPCs. To validate MIEL’s ability to discriminate between different cellular states/fates involving major changes in chromatin organization (e.g., reprogramming and differentiation), we analyzed 3 cell types: primary human fibroblasts isolated from 3 donors (WT-61, WT-101, WT-126), induced pluripotent stem cell (iPSC) lines derived from the fibroblasts, and neural progenitor cell (NPC) lines differentiated from the iPSCs, therefore providing genetically matching fibroblasts, iPSC and NPC cells (the cell lines were kindly provided by the Moutri group, UCSD). Cellular identities of the 3 cell types were verified by immunofluorescence (Fig. 2b).

The 9 cell lines were immunostained for H3K4me1 and H3K9me3 marks, chosen based on major pattern alteration of these marks during differentiation (35,36). Note that immunostaining for H3K27ac and H3K27me3 marks produced a similar distance map (Supplementary Fig. 5a). Both pairs of epigenetic marks were used interchangeably for further analysis. We segmented images and extracted image features, as previously described (26). Multivariate centroids were calculated for each cell population. Multi-dimension scaling (MDS) was employed to reduce 524 texture features into 2D and plotted to visualize the relative Euclidean distance between various cell populations (referred to as the “distance map”). Fibroblasts, iPSCs and NPCs each segregate to form 3 visually distinct territories (Fig. 2c).

To determine whether it was possible to discriminate between individual cells with different fates, a Support Vector Machine (SVM) classifier was trained using fibroblasts, iPSCs, and NPCs derived from donor WT-61. This classifier accurately identified 79% of fibroblasts, 79% of iPSCs and 97% of NPCs (overall accuracy 85% Fig. 2d; overall accuracy for H3K27ac and H3K27me3 based classification was 82% Supplementary Fig. 5b). Similar results were obtained when the classifier was trained using cell lines from the other 2 donors. A classifier derived by pooling WT-61, WT-101, and WT-126 cells correctly identified 89% of fibroblasts, 90% of iPSCs and 94% of NPCs (overall accuracy 91% Fig. 2e; overall accuracy for H3K27ac and H3K27me3 based classification was 90% Supplementary Fig. 5c). Furthermore, a direct pairwise classification distinguished different genetic backgrounds with 74% (Supplementary Fig. 5d). Additionally, MIEL analysis was able to discriminate between various primary hematopoietic cell types freshly isolated from mouse bone marrow suggesting that such analysis is not a cell culture artifact (Supplementary Fig. 6).

These results suggest that MIEL can be used to distinguish between different differentiation states based on their single-cell epigenetic landscapes. Furthermore, we were able to derive multiparametric signatures for several cell types (e.g., fibroblasts, iPSCs, NPCs) that discriminate each cell type from the others.

### MIEL determines signature of TPC differentiation

To begin deriving the signature of GBM differentiation, we tested MIEL’s ability to distinguish TPCs and differentiated glioma cells (DGCs), derived from the same primary human GBMs (6). Three TPC/DGC pairs (kindly provided by the Bernstein group, MGH, Harvard) were derived in parallel from 3 genetically distinct GBM tumor samples (MGG4, MGG6, and MGG8) over a 3-month period using either serum-free FGF/EGF conditions for TPCs or 10% serum for DGCs (6). MIEL analysis distinguished TPCs from their corresponding DGC lines with an average accuracy of 83%, using any of the 4 epigenetic marks tested (H3K27me3, H3K9me3, H3K27ac, and H3K4me1; Fig. 3a, b). An SVM classifier derived from images of the MGG4 TPC/DGC pair separated all 3 TPC/DGC pairs with 88% average accuracy, providing proof of principle for the derivation of a signature for non-tumorigenic cells obtained following serum differentiation of primary GBM cells (Fig. 3c).

**Figure 3.**
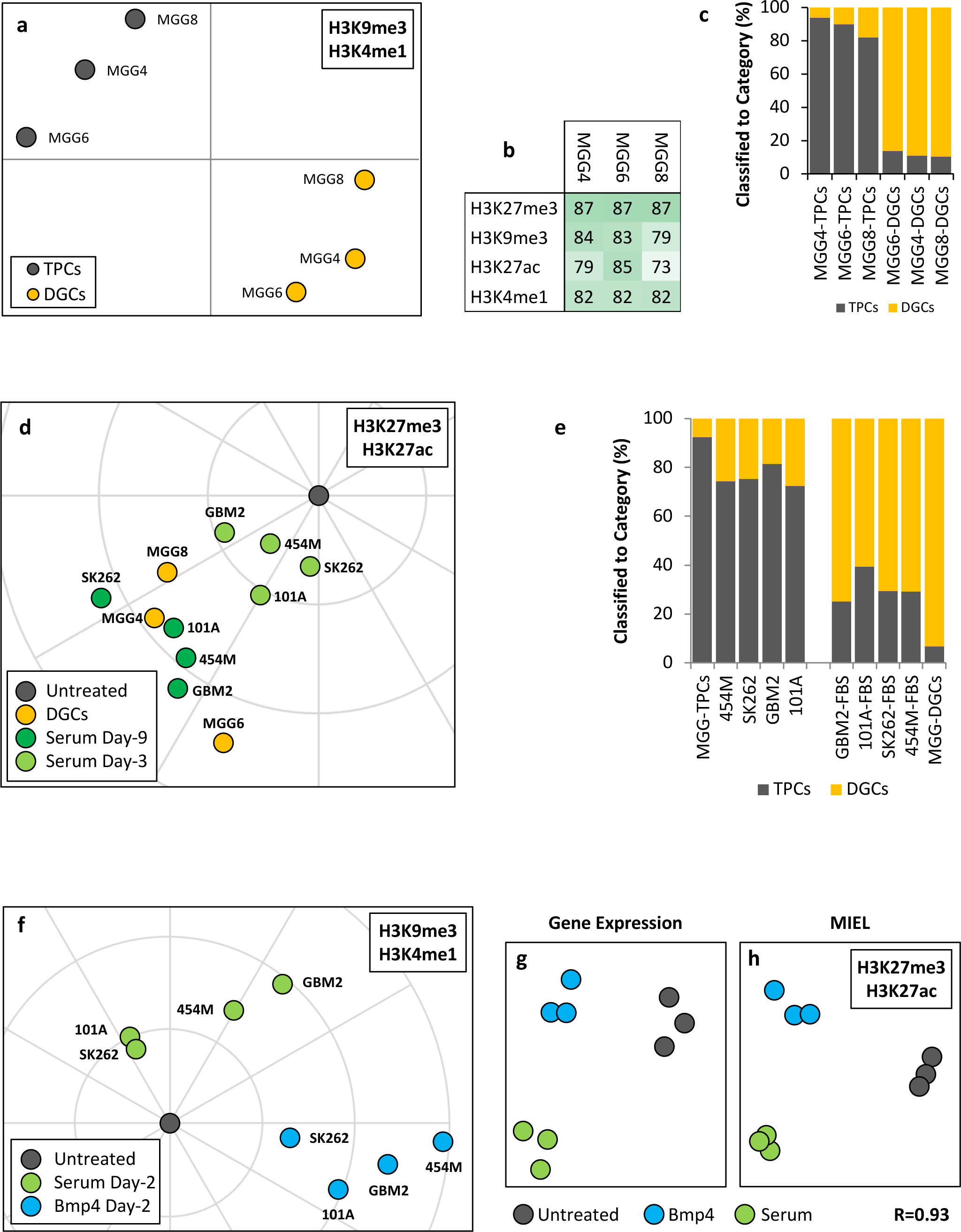
MIEL determines signature of TPC differentiation. (a) Distance map depicting the relative Euclidean distance between the multiparametric centroids of 3 genetically distinct TCP and DGC lines calculated using texture features derived from images of H3K9me3 and H3K4me1 marks. (b) TPC and DGC cell lines derived from the same tumor were distinguished by MIEL using multiple epigenetic marks. Numbers (percentages) correspond to the accuracy of TPC vs DGC pairwise classifications for each line using the indicated epigenetic marks. (c) Classification of TPC and DGC lines using an SVM classifier trained on texture features derived from images of H3K27ac and H3K27me3 marks in the MGG4 line. (d) Polar plot visualizing changes in feature values derived from H3K27ac and H3K27me3 images of 4 genetically distinct GBM line (GBM2, 454M, 101A, SK262) treated with serum for 3 or 9 days as well as TPC-DGCs pairs from MGG4, 6, 8 lines. (e) Two-way classification of 9 days serum-treated GBM lines, TPC, and DGCs from pooled MGG4, 6, 8 lines (MGG-TPC and MGG-DGC) using an SVM classifier trained on texture features derived from MGG-TPC and MGG-DGC. (f) Polar plot visualizing changes in feature values derived from H3K9me3 and H3K4me1 images of 4 GBM lines (GBM2, 454M, 101A, SK262) treated for 2 days with either serum or Bmp4. Average values for all four lines: FBS-2d: rho=5.7±2.7, theta=1.7±0.7; BMP4-2d: rho=9.6±3.8; theta=-0.2±0.1. (g) Distance map depicting the relative Euclidean distance between the transcriptomic profiles of DMSO-, Bmp4- and serum-treated GBM2 cells calculated using FPKM values of all expressed genes (14,376 genes; FPKM>1 in at least one sample). Each treatment is in triplicates. (h) Distance map depicting the relative Euclidean distance between the multiparametric centroids of DMSO-, Bmp4- and serum-treated GBM2 cells calculated using texture features derived from images of H3K27ac and H3K27me3 marks. Each treatment is in triplicates. R denotes Pearson correlation coefficient.

Next, we asked whether shorter serum treatment (compatible with the screening protocols) would induce detectable epigenetic alterations. We treated 4 low-passage primary TPCs for 9 days with 10% serum and compared their epigenetic landscape to that of untreated cells and “terminally” differentiated DGCs. As before, we used MDS to visualize the relative Euclidean distance between populations. While untreated cells were quite heterogeneous, serum treatment reduced the distance from all TPC centroids to DGC centroids (n=4 cell lines, p<0.05; unpaired two-tailed t-test; Supplementary Fig. 7a, b). These concordant results obtained with 7 independent human GBM lines attest to the robustness of the serum-induced epigenetic changes detected by MIEL and suggesting that similar epigenetic patterns may exist in other differentiated GBM lines.

To compare the outcomes of several experiments using multiple GBM lines and treatments, we developed a normalization procedure that compares the changes in feature space induced by treatments by bringing together the centroids of all TPCs (including MGG-TPCs). The results are then displayed using a polar plot in which treatments for each cell line are represented as vectors with a magnitude - rho (the distance from the center) and directionality given by the angular coordinate theta. Remarkably, for all GBM lines, the magnitude and direction of changes induced by 9-day serum treatment were comparable and similar to that seen in the MGG TPC/DGC pairs (rho: FBS-9d=10.1±1.0, DGCs=10.0±1.64; theta: FBS-9d=-2.4±0.2, DGCs=-2.5±0.4); three-day serum treatment induced feature changes comparable in direction, but not magnitude (rho: FBS-3d=4.1±1.0; theta: FBS-3d=-2.2±0.5; Fig. 3d and Supplementary Fig. 7c).

To test the accuracy of separating TPCs and DGCs at the single-cell level we generated an SVM classifier trained on texture features derived from a random subset of H3K27ac and H3K27me3 images of TPCs and DGCs (MGG4,6,8 pooled for both). The classifier separated pooled TPCs from pooled DGCs with 92.8% accuracy (Fig. 3e) and categorized 76% of untreated cells as TPCs and 69% of serum-treated cells as DGCs (Fig. 3e).

These experiments demonstrate that MIEL is suitable to determine a signature of differentiated GBM cells across multiple genetic backgrounds. Furthermore, MIEL can detect serum-induced changes in GBM epigenetic pattern within several days to monitor the progress of TPC differentiation in a timeframe suitable for high content screening.

### Validation of MIEL signature using global transcriptomic analysis

Previous work indicated distinct features of GBM differentiation induced with BMP compared to serum (19). Indeed, we observed distinct expression changes, including differences in expression of genes regulating chromatin organization and histone modifications (Supplementary Fig. 8), between serum- and Bmp4-induced GBM differentiation. Therefore, we inquired whether MIEL approach could distinguish these differentiation modalities, in particular at the early time points.

We treated 4 genetically distinct GBM lines for 2 days with serum or BMP4 and conducted MIEL analysis using H3K9me3 and H3K4me1 marks. To visualize the changes induced by each treatment, we used polar plot normalization, as described above. Indeed, we observed that serum and BMP4 induce distinct epigenetic changes as detected by MIEL for each GBM line tested (Fig. 3f).

Global gene expression profile represents a gold standard to define the cellular state (37). Therefore, we asked whether the relative distances between distinct cellular states, for instance, untreated GBM cells, serum treated, and BMP treated GBM cells correlate using MIEL-based metrics and global gene expression-based metrics. We sequenced untreated and 3 days serum or Bmp4 treated GBM2 TPCs. All genes with FPKM>1 in at least one cell population were used to calculate the Euclidean distance matrix between all cell populations. FPKM-based distances were then correlated to image texture feature-based distances. The resulting Pearson correlation coefficient of R=0.93 suggests a high correlation between these 2 metrics (Fig. 3g, h) and validates the robustness of MIEL approach for the analysis of GBM differentiation.

These experiments demonstrate that MILE is capable of distinguishing closely related GBM differentiation routes induced by serum or BMP. Critically, these results validate the robustness and accuracy of MIEL-based analysis of epigenetic patterns using conventional global gene expression approach.

### MIEL prioritizes compounds based on serum/Bmp4 signature of GBM differentiation

To test whether MIEL can prioritize compounds based on serum/Bmp4 signature of GBM differentiation, we re-screened the Prestwick compound library (at lower concentration, 3 μM to minimize toxicity). GBM2 TPCs were plated on 384-well plates, treated for 3 days with Prestwick compounds fixed, and then immunostained for H3K27ac and H3K27me3. GBM2 cells treated with DMSO, serum, BMP4, or compound were compared within the same plate (to avoid imaging artifacts and normalization issues). To identify compounds inducing epigenetic changes reminiscent of serum/BMP4-induced differentiation, we conducted pairwise classification of DMSO- and either serum- or BMP4-treated cells. Because both serum and BMP4 induce TPC differentiation and reduce tumorigenicity we selected compounds that induced at least 50% of the cells to be classified as either serum- or BMP4-treated. We then calculated the Euclidean distance between these candidate compounds and serum/BMP4 treated cells – selecting compounds for which the distance to one or both treatments was less than the distance between DMSO and that treatment. This screen yielded 20 candidate compounds (Supplementary Fig. 9a), of which 15 belonged to 1 of the following 4 categories: Na/K-ATPase inhibitors of the digoxin family, molecules that disrupt microtubule formation or stability, topoisomerase inhibitors, and nucleotide analogues that disrupt DNA synthesis.

Of these 15 candidate compounds, we chose 2 top compounds from each of the 4 categories (8 total) for further analysis. For each of the 8 compounds, we used pairwise classification of untreated cells and either serum- or Bmp4-treated cells to identify the lowest concentration where at least 50% of cells are categorized as treated (Supplementary Fig. 9b). These concentrations were used for all subsequent experiments. Because most of these compounds are known for their cytotoxic effects, we verified the growth rates of drug-treated GBM cells. With the exception of Digoxin, which was cytostatic, treatment with drugs resulted in the growth rates comparable with that induced by serum/BMP4 treatment (supplementary Fig. 10a). We used immunofluorescence to test for the expression of the core TPC transcription factors (Sox2, Sall2, Brn2 and Olig2). With the exception of Trifluridine all compounds induced statistically significant reductions in Sox2, but no reduction in the other core factors (Supplementary Fig. 10b; see Fig. 1c for Digoxin and Digitoxigenin)

Next, we investigated whether MIEL can prioritize compounds according to their effect on TPC as judged by the transcriptomic changes induced by these compounds. GBM2 cells were treated with DMSO (negative control), serum or Bmp4 (positive controls), or 1 of the 8 candidate compounds; after 3 days, RNA was extracted and sequenced. Transcriptomic profiles of the 8 compounds were ranked according to average Euclidean distance (based on FPKM values for all expressed genes) from serum/BMP4-treated cells. To safeguard against potential artefacts of cytotoxicity we compared gene expression-based ranking with the measured cellular growth rates for all drug treatments. Indeed, no positive correlation was revealed (Supplementary Fig. 10c).

Next, we compared the levels of Sox2 expression under all treatment conditions to determine whether this metric is informative for identifying the drugs that best mimic serum/BMP4 treatment. We did not observe a positive correlation between Sox2 expression levels and the transcriptomic-based rankings (Fig. 4a), suggesting that SOX2 level alone is insufficient to stratify the compounds.

**Figure 4.**
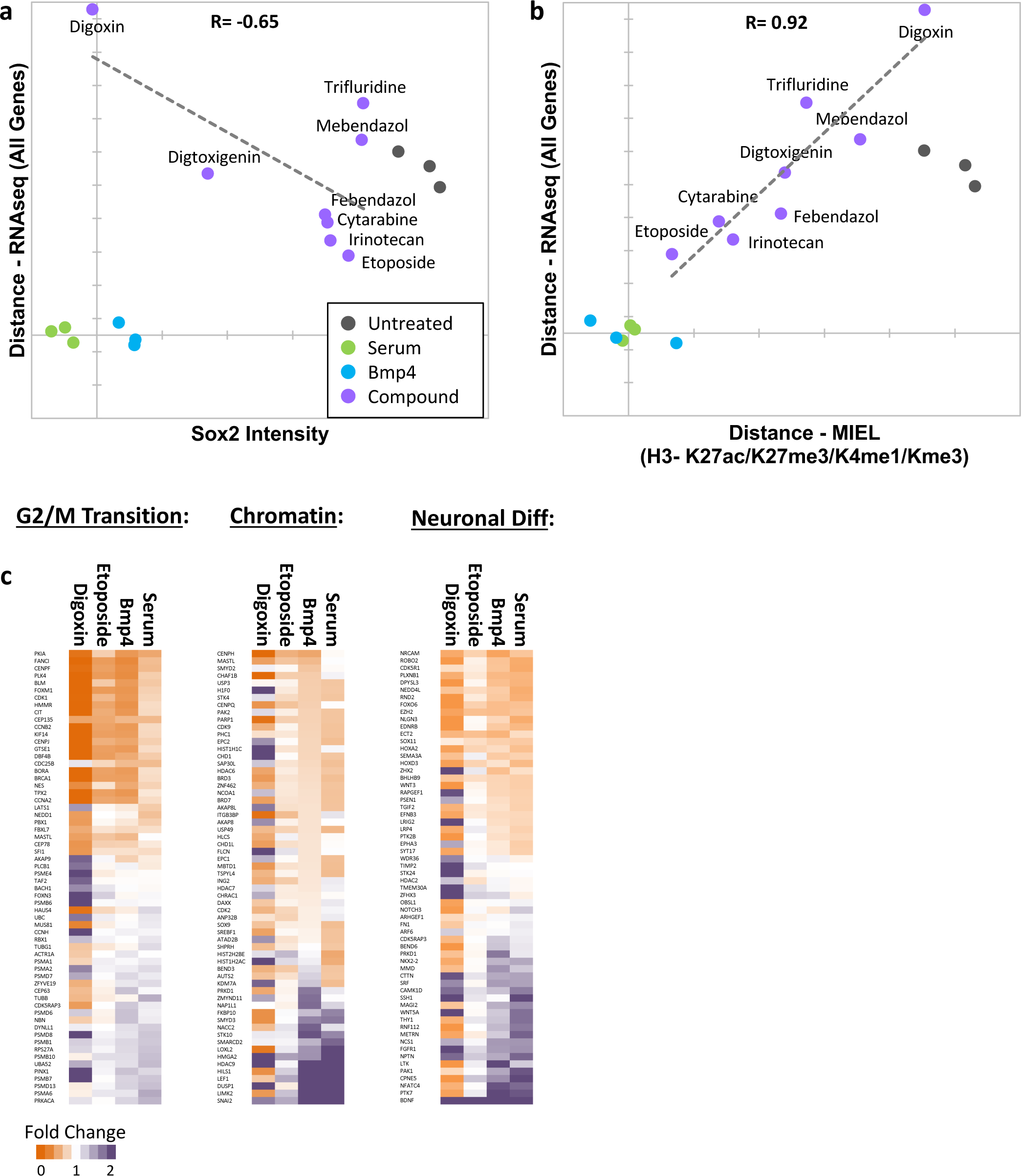
MIEL prioritizes small molecules based on serum/Bmp4 differentiation signature. (a) Scatter plot showing the correlation of gene expression profile-based ranking and Sox2 expression for 8 candidate drugs, untreated, serum or Bmp4 treated GBM2 cells. Euclidean distance to serum or Bmp4 treated GBM2 cells was calculated using transcriptomic profiles (vertical axis), or Sox2 immunofluorescence intensity (horizontal axis). Distances and Sox2 levels were normalized to untreated and serum/Bmp4 treated GBM2 cells. (b) Scatter plot showing the correlation of gene expression profile-based ranking and MIEL-based ranking for 8 candidate drugs, untreated, serum or Bmp4 treated GBM2 cells. Euclidean distance to serum- or Bmp4-treated GBM2 cells was calculated using transcriptomic profiles (vertical axis), or texture features derived from images of H3K27ac and H3K27me3, H3K9me3, and H3K4me1 marks (horizontal axis). Distances were normalized to untreated and serum/Bmp4 treated GBM2 cells. (c) Heat maps showing fold change in expression of select genes taken from Gene Ontology list: cell cycle G2/M phase transition (GO:0044839), chromatin modification (GO:0006325), and regulation of neuron differentiation (GO:0045664). R denotes Pearson correlation coefficient. Drug concentrations a-c: febendazole=0.5 μM, mebendazole=0.5 μM, cytarabine=0.3 μM, trifluridine=3 μM, irinotecan=0.5 μM, etoposide=0.3 μM, digitoxigenin=0.3 μM, digoxin=0.3 μM.

To compare MIEL based signatures to the transcriptomic profile we first sought to get a comprehensive readout of the epigenetic landscape of treated cells. We therefore conducted MIEL analysis using an additional set of histone modifications including H3K9me3 and H3K4me1 marks. We then ranked MIEL readouts of cells treated with the 8 drugs according to average Euclidean distance from serum- or Bmp4-treated cells (calculated using texture features derived from images of 4 histone modifications). Comparison of the MIEL-based metric with the gene expression-based metric revealed a high degree of positive correlation between MIEL- and gene expression-based rankings (Pearson correlation coefficient R=0.92, p<0.001, one side t-test, n=6, Fig. 4b). To further visualize these results, we constructed heat-maps depicting fold change in expression of genes associated with several GO terms enriched by serum and Bmp4 treatments. Our top candidate, etoposide, altered expression of a large portion of genes in a similar fashion to that of serum and BMP4; in contrast, the lowest-ranking candidate, digoxin, induced gene expression changes that were rather different from serum and BMP4 (Fig. 4c).

The above results suggest a robust correlation between MIEL- and global expression-based readouts of GBM differentiation, therefore validating MIEL approach for prioritizing hits in high-content screening aimed at identifying small molecules that mimic the effect of serum/BMP4 on GBM differentiation.

## Discussion

Cytotoxic drugs have had limited success treating GBM; therefore, we focused on the alternative approach - inducing GBM differentiation. We analyzed previously established biologicals such as serum and BMP4 known to induce GBM differentiation in culture (16–18) and established signatures of such differentiated GBM cells based on the pattern of epigenetic marks that could be applied across several genetic backgrounds. This is the first time that GBM differentiation signature suitable for high-throughput drug screening could be obtained. Indeed, the results of previous studies using bulk analysis of GBM (19) or single-cell sequencing (23) could not be readily applied for high-throughput screening. As a proof of principle, we analyzed Prestwick chemical library of 1200 approved drugs to validate MIEL’s ability to select and prioritize small molecules, which mimic the effect of serum and BMP4 using global gene expression profiling. Surprisingly, we observed that the degree of reduction in endogenous SOX2 protein levels following drug treatment did not correlate with the degree of differentiation assessed by global gene expression. In contrast, the MIEL-based metrics did correlate with the degree of differentiation assessed by global gene expression. Therefore, MIEL can be readily applied to screen large compound libraries using a reference signatures of GBM differentiation (e.g. serum or BMP4) to identify novel small molecules that mimic the effect of serum or BMP4 on GBM.

Accurately defining the identity of a cell is of fundamental importance to cell biology. Currently, this is done by assessing the presence or absence of a panel of experimentally verified lineage-specific markers. However, these markers require manual and arbitrary thresholding, which could be confusing and potentially contributes to multiple challenges of reproducibility in biomedical science (38,39). These concerns are alleviated by expression profiles that use hundreds of genes to assign a specific gene signature to a given cell type. However, at a single-cell level, expression profiling becomes stochastic, and is difficult to apply to high-throughput analysis in a cost- and time-effective manner. Phenotypic drug screening is an emerging technology that is revolutionizing drug discovery (26,33,40). Here, we described a new method for phenotypic identification of a cell state that offers reproducibility, single-cell resolution and scalability for high-content screening. MIEL takes advantage of robust and reproducible patterns of epigenetic marks that are always present in every eukaryotic cell. As a proof of principle using MIEL, we were able to define the unique signatures of various cell types in culture such as fibroblasts, iPSCs, NPCs as well as primary cells isolated from mouse bone marrow (T cells, B cells, monocytes, and hematopoietic stem/progenitor cells) enabling their identification with over 80% accuracy.

It is becoming increasingly apparent that nuclear chromatin is spatially organized relative to the gene expression pattern (41); for example, CTCF proteins play a role in dictating boundaries of topologically associated domains (TADs) (42,43). TADs are thought to parse chromatin into loosely defined active (euchromatin) and inactive (heterochromatin) domains, reciprocating particular patterns of gene activity (41). It is tempting to conjecture that the 2D epigenetic landscapes, which can be imaged at the single-cell level by MIEL, define the state of chromatin and the gene expression pattern (ie, a cell’s molecular identity). While other phenotypic screens are based on diverse strategies for labeling cellular compartments (e.g. nucleus, membranes or mitochondria) (33,40), MIEL is rooted in the spatial organization of epigenetic marks. The ability of MIEL to distinguish between multiple cell fates with high accuracy indicates that the topology of epigenetic marks might be used as a proxy for single cell state and function. Providing MIEL can be adapted to analyze epigenetic landscapes in 3D, it might offer unique insights into cellular heterogeneity during development and aging and enable in situ analysis of epigenetic variations in normal human tissues and various pathologies including cancer.

## Materials and Methods

### Cell Culture

Monolayer cultures of patient-derived TPCs were propagated on Matrigel-coated plates in DMEM:F12 Neurobasal media (1:1; Gibco), 1% B27 supplement (Gibco), 10% BIT 9500 (StemCell Technologies), 1 mM glutamine, 20 ng/ml EGF (Chemicon), 20 ng/ml bFGF, 5 μg/ml insulin (Sigma), and 5 mM nicotinamide (Sigma). The medium was replaced every other day and the cells were enzymatically dissociated using Accutase prior to splitting. Fibroblasts, iPSCs, and iPSC-derived NPCs were cultured as previously described(44,45).

### Differentiation treatment

For TPC differentiation treatments cells were cultured in DMEM:F12 Neurobasal media (1:1), 1% B27 supplement, 10% BIT 9500, 1 mM glutamine supplemented with either Bmp4 (100ng/ml; R&D Systems) or FBS (10%).

### Cell staining

Cells were rinsed with PBS and fixed in 4% paraformaldehyde in PBS for 10 min at room temperature. After blocking with PBSAT (2% BSA and 0.5% Triton X-100 in PBS) for 1 h at room temperature, the cells were incubated overnight at 4°C with primary antibodies diluted in PBSAT. The primary antibodies are listed in Table 2, and the appropriate fluorochrome-conjugated secondary antibodies were used at 1:500 dilution. Nuclear co-staining was performed by incubating cells with Hoechst-33342 nuclear dye.

### RNAseq and transcriptomic analysis

Total RNA was isolated from GBM2 cells using the RNeasy Kit (Qiagen), 0.5 ug total RNA was used for isolation of mRNAs and library preparation. Library preparation and sequencing was conducted by the SBP genomics core (Sanford-Burnham NCI Cancer Center Support Grant P30 CA030199). PolyA RNA was isolated using the NEBNext^®^ Poly(A) mRNA Magnetic Isolation Module and barcoded libraries were made using the NEBNext^®^ Ultra II^™^ Directional RNA Library Prep Kit for Illumina^®^(NEB, Ipswich MA). Libraries were pooled and single end sequenced (1X75) on the Illumina NextSeq 500 using the High output V2 kit (Illumina). Read data was processed in BaseSpace (basespace.illumina.com). Reads were aligned to Homo sapiens genome (hg19) using STAR aligner (https://code.google.com/p/rna-star/) with default settings. Differential transcript expression was determined using the Cufflinks Cuffdiff package (https://github.com/cole-trapnell-lab/cufflinks). Go term enrichment analysis was conducted using PANTHER v11 using all genes identified as differentially expressed following either serum or Bmp4 treatment. For heat maps showing fold change in expression the FPKM values in each population were divided by the average FPKM values of untreated GBM2. To highlight differences in expression levels between serum and Bmp4 treated GBM2 cells the FPKM values in each sample were z-scored. Zscore=(FPKM_Observation_-FPKM_Average_)/FPKM_SD_ (FPKM_Observation_-FPKM value obtain through sequencing; FPKM_Average_ – average of all FPKM values in all samples for a certain gene; FPKM_SD_ – standard deviation of FPKM values for a certain gene). Heat maps were generated using Microsoft Excel conditional formatting function.

### Prestwick Chemical Library screen using Sox2 and GFAP

GBM2 cells were plated at 2000 cells/well and exposed to Prestwick compounds (10 μM) for 3 days in 384-well optical bottom assay plates (Greiner). Cells were then fixed and stained with goat polyclonal anti-Sox2 and rabbit polyclonal anti-GFAP (Table 2) antibodies followed by AlexaFluor-488- or AlexaFluor-555-conjugated secondary antibodies. The positive and negative control treatments were BMP4 (100 ng/ml) and DMSO (0.1%), respectively. DNA was counterstained with DAPI and the cytoplasmic region was identified with HCS CellMask Deep Red. Images were acquired using the Perkin Elmer Opera^®^ QEHS. Image analysis protocols were developed with PerkinElmer Acapella^®^ using standardized analysis building blocks and custom algorithm scripting. Specific antibody-based parameters, morphological and fluorometric parameters, and nuclei counts were extracted for the imaged region in each well. Nuclear mask was segmented based on DAPI stain, cytoplasm mask was segmented based on CellMask. Image analysis included quantification of cell count, the nuclear staining intensity of Sox2 and the cytoplasmic intensity of GFAP. These parameters were used to evaluate activity of compounds, which was scored as percent efficacy for decrease in Sox2 levels and increase in GFAP levels. The average robust Z’-scores (RZ’) is based on the Z′-score (46) but uses the median and the median absolute deviation instead of the mean and the standard deviation. RZ’ were 0.31 and 0.29 for Sox2 and GFAP, respectively. Percent efficacy was calculated as: Percent efficacy=((Obs-NegCont)/(PosCont-NegCont))X100; Obs, intensity measured for compound; NegCont, average intensity of 32 DMSO-treated wells in each plate; PosCont - average intensity of 32 Bmp4-treated wells in each plate. Percent efficacy for each compound was calculated using only controls from the same plate. Hits were defined as compounds that yield percent efficacy values of either GFAP (increase) and Sox2 (decrease) >40, or only Sox2 decrease >100. To evaluate drug induced cytotoxicity robust z-score for the number of non-Pyknotic cells (Pyknotic cells were identified by decreased nuclear area and increased DAPI intensity) was calculated according to: RZscore=(Count_Observation_-Count_Median_)/Count_MAD_ where Count_Observation_ denotes count of viable nuclei in the well; Count_Median_ denotes median cell count for all DMSO treated wells; Count_MAD_ denotes median absolute difference of cell count for all DMSO treated wells).

### Microscopy and image analysis

Unless stated otherwise, for MIEL analysis cells were imaged on an Opera QEHS high-content screening system (PerkinElmer) using ×40 water immersion objectives. Images collected on the Opera were analyzed using Acapella 2.6 (PerkinElmer). At least 40 fields per well were acquired and at least 2 wells per population were used. Features of nuclear morphology, fluorescence intensity inter-channel co-localization, and texture features (Image moments, Haralick, Threshold Adjacency Statistics) were calculated using custom algorithms (scripts available from www.andrewslab.ca). A full list of the features used is available from the authors. Values for each cell were generated and exported to MATLAB for further analysis. For Sall2, Olig2, Brn2, Sox2, Oct4 and GFAP immunostaining, images were captured on an IC200-KIC (Vala Sciences) using a ×20 objective. Between 3 and 8 fields per well were acquired and analyzed using Acapella 2.6 (PerkinElmer). For all nuclear markers, average intensities in nucleus or fold change in average intensity compared to untreated cells are shown. Unless stated otherwise, at least 3 wells and a minimum of 300 cells for each condition were compared using unpaired two-tailed t-test was.

### MDS

The image features based profile for each cell population (eg, cell types, treatments) was represented using a vector (center of distribution vectors) in which every element is the average value of all cells in that population for a particular feature. The vector’s length is given by the number of features chosen. All vectors used to composite the MDS maps (distance maps) consisted of 524 texture features (262 per channel, 2 channels). Cell-level data in all populations together were normalized to z-scores prior to calculation of center of distribution vectors. All cells in each population were used to calculate center vectors and each population contained at least 400 cells. Transcriptomic based profile for each cell population was represented using a vector in which every element is the z-scored FPKM value for a single gene in that population. The length of the vector is given by the number of genes used to construct the profile. The Euclidean distance between all vectors (either image features or transcriptomic based) was then calculated to assemble a dissimilarity matrix (size N×N, where N is the number of populations being compared). For representation, the N×N matrix was reduced to a Nx2 matrix with MDS using the MATLAB (2016a) function ‘cmdscale’ or an Excel add-on program Xlstat (Base, v19.06), and displayed as a 2D scatter plot.

### Polar plots

Due to the inherent heterogeneity of TPC lines, we performed data normalization when comparing multiple treatments on several TPC lines. For this, the value of each feature for all individual cells in each line was divided by the average value obtained for that feature in the untreated population from the same cell line. Therefore, following normalization, untreated cells from all lines had the same center of distribution vector (in which all elements are equal to 1), while each treatment retained its relative distance from untreated as well as from all other treatments of the same cell line. However, as each cell line is divided by a different value, the distance vectors originating from two different lines represent the change in feature values induced by treatment, rather than the absolute feature values. Therefore, following MDS, the results are shown on a polar plot to indicate that the various treatments induce similar feature value changes in multiple lines rather than similar absolute values. As a result, direction and distance to the origin are comparable between lines while distances directly between points are not.

### SVM classification

SVM classification was conducted as previously described (26). Cell-level data in all populations (minimum 400 cells per population) together were normalized to z-scores and a subset of cells from each of the populations being classified was randomly chosen as the training set (subset size is at least 100× the number of populations being classified). The training set was used to train a SVM classifier (MATLAB function ‘svmtrain’). The remaining cells (test set) were then classified using the SVM-derived classifier to assess the accuracy of classification (MATLAB function ‘svmclassify’). Here, the accuracy of all pairwise classifications is given as the average accuracy calculated for each of the populations. To utilize classification to determine the similarity of multiple cell populations, we classified known populations (such as different treatments or cell fates) to generate known ‘bins’ and then used the same classifiers on the unknown population to categorize each cell.

### Prestwick Chemical Library screen using H3K27me3 and H3K27ac

GBM2 cells were plated at 2000 cells/well and exposed to Prestwick compounds (3 μM) for 3 days in 384-well optical bottom assay plates (PerkinElmer). Cells were then fixed and stained with rabbit polyclonal anti-H3K27ac and mouse monoclonal anti-H3K27me3 (Table 2) antibodies followed by AlexaFluor-488- or AlexaFluor-555-conjugated secondary antibodies. The positive control treatments were BMP4 (100 ng/ml) and serum (10%), negative controls were DMSO (0.1%). DNA was counterstained with Hoechst. Images were acquired using the Perkin Elmer Opera^®^ QEHS. MIEL analysis was conducted as described above. The robust Z′-score (RZ’) is based on the Z′-score described in (32), but uses the median (<*x*>) and the robust standard deviation (rSD) based on the median absolute deviation (MAD) instead of the mean and the standard deviation. Briefly, we use the DMSO-(negative) and BMP4- or serum-treated (positive) control wells to establish the signatures corresponding to undifferentiated (DMSO) and differentiated (BMP4 and/or serum) GBM cells. Using these signatures, we then classify the cells in each well to obtain the population fraction of differentiated GBM cells per well. These values are used to calculate the medians and rSDs for all DMSO (<*x*>_neg_ and rSD_neg_) and all BMP- or serum-treated (<*x*>_pos_ and rSD_pos_) wells. The RZ’ value is calculated as follows: RZ′=1 - (3*rSD_pos_+3*rSD_neg_)/|<*x*>_pos_-<*x*>_neg_| with rSD=MAD*1.4826 and MAD=<|*x* -<*x*>|>. RZ’ values are calculated for DMSO vs. Bmp4 treated, DMSO vs. serum treated, and DMSO vs. pooled Bmp4 and Serum treated wells. The Signal-to-Background (S/B) described in (32) uses the formula: μ_pos_ / μ_neg_ where μ is the average of the differentiated population fractions for all treated (Bmp4, Serum) or control (DMSO) wells.

### Correlation of transcriptomic and image-based profiles

Euclidean distance between untreated, serum or Bmp4 treated GBM2 cells (triplicates for each) was calculated using either transcriptomic data (FPKM) or texture features. Pearson’s correlation coefficient (R) was transformed to a t-value using the formula (t = R × SQRT(N-2)/SQRT(1-R2) where N is the number of samples, and R is Pearson correlation coefficient, and the p-value was calculated using Excel tdist(t) function. For compound prioritization, the Euclidean distance between compound treated and serum or Bmp4 treated GBM2 cells was calculated based on either transcriptomic data (FPKM) or image features. The average distance to both serum and Bmp4 treatments was normalized to the average distance of untreated cells to serum and Bmp4.

## Acknowledgments

We are thankful to Laure Escoubet (Celgene) for discussions and support along the entire project, Alex Kiselyov (Genea Biocells) for his invaluable help with compound libraries and discussions, Harley Kornblum (UCLA) for sharing multiple primary human GBM lines, to Alysson Muotri (UCSD) for providing fibroblast, iPSC and NPC lines, and Bradley Bernstein (MGH Harvard) for sharing MGG-TPCs and MGG-DGCs lines. We owe a debt of gratitude to Susanne Heynen-Genel, Debbie Chen, and other members of the High-Content Facility at CPCCG for their invaluable help with cell imaging and to Brian James and Kang Liu at the SBP Genomics core for their help with library preparation and RNA sequencing (NCI Cancer Center Support Grant P30 CA030199). We thank Linda Penn for suggestions and help with the manuscript. This work was supported by sponsored research agreement with Celgene and an R01 NS066278 to A.V.T and by a CIHR Foundation grant and a Tier 1 Canada Research Chair award to D.W.A.

Author contributions
C.F. developed novel approaches and analyzed the data, wrote the manuscript, and contributed to all aspects of this study, C.F., L.V.W, F. C., M. N., performed the experiments, S. H., J. Y., wrote the code and helped with the data analysis, D.W.A and A.V.T. conceived the idea, wrote the manuscript, and guided all aspects of this study.

